# Polygenic scores for UK Biobank scale data

**DOI:** 10.1101/252270

**Authors:** Timothy Shin Heng Mak, Robert Milan Porsch, Shing Wan Choi, Pak Chung Sham

## Abstract

Polygenic scores (PGS) are estimated scores representing the genetic tendency of an individual for a disease or trait and have become an indispensible tool in a variety of analyses. Typically they are linear combination of the genotypes of a large number of SNPs, with the weights calculated from an external source, such as summary statistics from large meta-analyses. Recently cohorts with genetic data have become very large, such that it would be a waste if the raw data were not made use of in constructing PGS. Making use of raw data in calculating PGS, however, presents us with problems of overfitting. Here we discuss the essence of overfitting as applied in PGS calculations and highlight the difference between overfitting due to the overlap between the target and the discovery data (OTD), and overfitting due to the overlap between the target the the validation data (OTV). We propose two methods — cross prediction and split validation — to overcome OTD and OTV respectively. Using these two methods, PGS can be calculated using raw data without overfitting. We show that PGSs thus calculated have better predictive power than those using summary statistics alone for six phenotypes in the UK Biobank data.

## Introduction

Polygenic scores, or polygenic risk scores (PGS), have become an indispensible tool in genetic studies [1, 2, 3, 4, 5, 6, 7, 8, 9, 10, 11, 12, 13, 14]. Polygenic scores are routinely calculated in small and large cohorts with genotype data, and they represent individual genetic tendencies for particular traits or diseases. As such they can be used for stratifying individuals into different risk groups based on their genetic makeup [14, 3, 4, 15]. Potentially, different interventions could be given to individuals with different risks, which is part of the vision in personalized medicine [16, 17].

Currently, however, the predictive ability of PGS for complex traits remains considerably lower than the maximum possible given their heritability, although with increasing sample sizes and the number of Genome-wide association studies, the power is set to increase [18, 19, 12]. Nonetheless, even before the objective of personalized medicine can be achieved, PGS can be used for studying the genetic influence of different phenotypes. By examining the correlation between PGS and various phenotypes, researchers can gather evidence for whether the genetic influence on certain traits were pleiotropic or specific [20, 21, 22, 23, 11, 6, 7]. For example, using PGS, Power *et al*[11] showed that genetic tendency for schizophrenia and bipolar disorder were predictive of creativity, supporting earlier suggestions that creativity and tendency towards major psychotic illnesses may share some common roots.

Polygenic scores are calculated as weighted sums of the genotypes, with weights typically derived from large cohorts or meta-analyses. A key requirement in the calculation of PGS is that the same individuals be not used both in the calculation of the weights (in the discovery dataset) and the PGS (in the target dataset). Indeed, in general, samples in the *discovery* and *target* dataset should not even be related [24]. Overlap or relatedness between the samples is expected to lead to *overfitting*, i.e. the inflation in measures of the fit in the target dataset.

Recently, cohorts with genotype data have become very large. Examples of such cohorts include the UK Biobank[25] (*n* ≈ 500,000), the 23andMe cohort [26] (*n* ≈ 600,000), and the deCode cohort [27] (*n* ≈ 350,000). In studies to date using the UK Biobank, for example, following the recommended practice, weights for the PGS were calculated from summary statistics and data external to the cohort [13, 28, 27]. Although sensible as a measure to avoid overfitting, the exclusion of the target dataset from the calculation of the summary statistics in these cases can be wasteful, given that such large sample sizes are involved.

In this paper, we show that it is possible to calculate PGS using the target dataset while avoiding overfitting, which can lead to higher predictive power than PGS calculated from summary statistics alone.

## Results

As an illustration of the potential gain in power using the target dataset in the calculation of PGS, consider the correlation between the phenotype and the PGS calculated using the method of this paper, which we call *cross prediction*, compared to using summary statistics only, as presented in Figure 1. The comparison is made using a cohort of 353,465 white British participants in the UK Biobank study [25]. We see that for all 6 phenotypes, using the data available in the UK Biobank alone gives a PGS with visibly higher correlation with the phenotype than the equivalent PGS calculated using summary statistics. The correlation was even higher when the UKBB data was meta-analysed with the summary statistics. In this section, we introduce the methods used in calculating the PGS in Figure 1. We show how these methods avoid *overfitting* and thus the improvement seen in Figure 1 is due to genuine increase in power because of the data available in the UK Biobank. We defer the details of the simulations to the Methods section at the end of the article.

**Figure 1:**
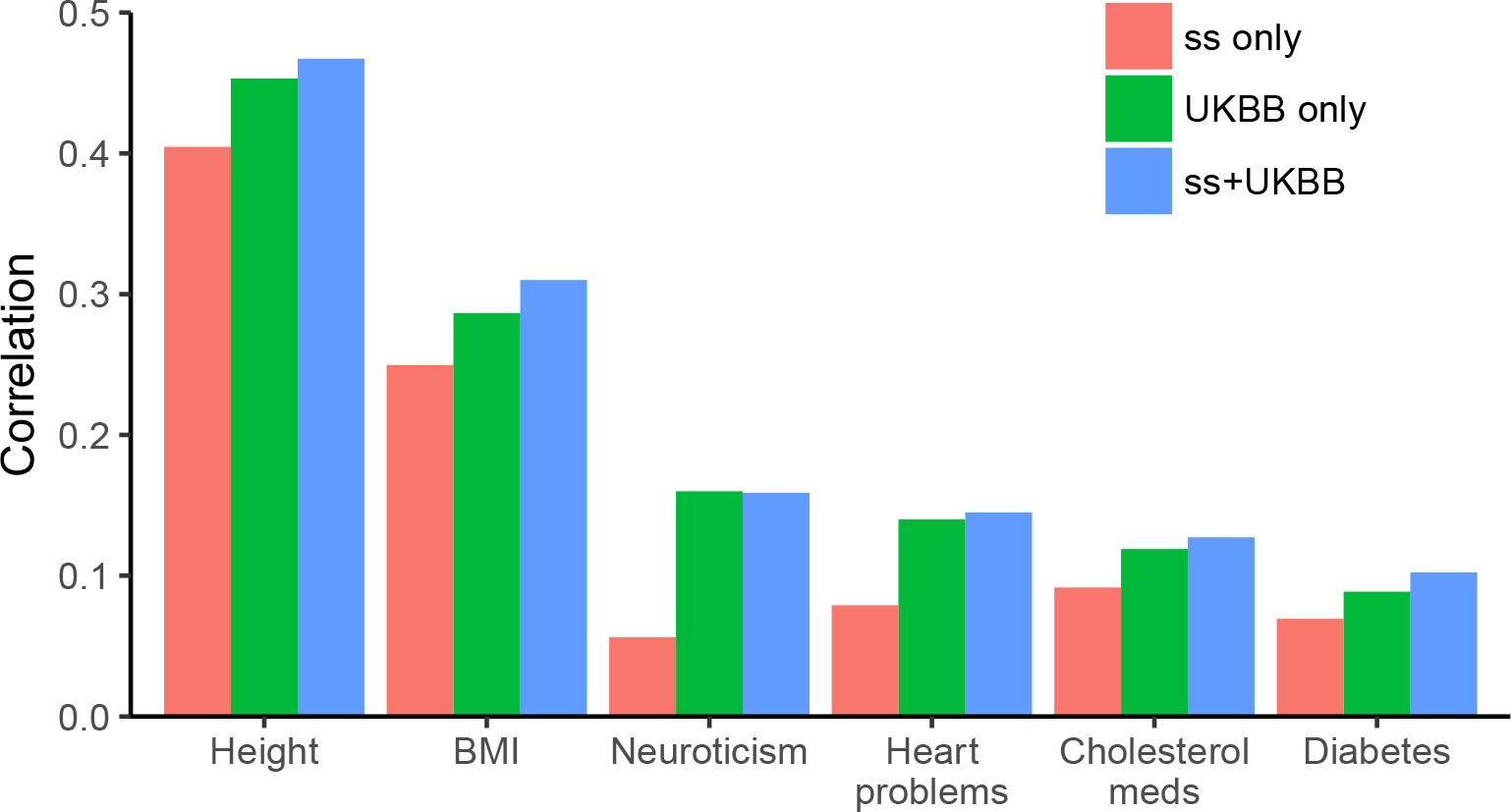
Correlation between phenotype and PGS calculated using summary statistics (ss) only, UKBB data only, and summary statistics plus UKBB data, in a cohort of 353,465 particpants in the UK Biobank.

### Three types of overfitting in calculating polygenic scores

In their review article, Wray *et al*[24] pointed out that if the same individuals were used in both the target dataset and the discovery dataset or if they were related, estimates of the predictive power of PGS would be inflated. Although not specifically mentioned, the phenomenon underlying this was that of *overfitting* of the data to the target dataset. Here, we define overfitting to be the inflation of the correlation of the PGS with the genetic component in the target dataset over a completely independent (unseen) external dataset. More precisely, let us assume the following linear model

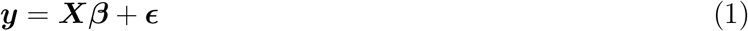

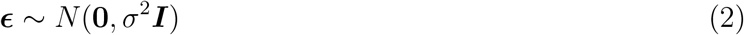

where **y** = (*y*_1_, *y*_2_,…, *y*_*n*_)′ denotes a vector of phenotype from *n* independent individuals from the *target* dataset. Let ***X β*** denote the genetic component and ***ϵ*** = (*ϵ*_1_, *ϵ*_2_,…, *ϵ*_*n*_)′ residual environmental effects, with *ϵ*_*i*_ assumed independently and identically distributed. We assume 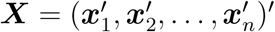 is a *n*-by-*p* genotype matrix and ***β*** a vector of causal effects. In the case where adjustment for principal components is necessary[29], we assume that both ***y*** and ***X*** have the principal components of ***X*** regressed out of them. A PGS for an individual *i* is an estimate of ***x***_*i*_***β***, denoted 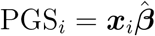. We define overfitting as

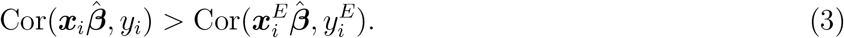

where (***x***_*i*_, *y_i_*) is a randomly chosen sample from the target dataset, and 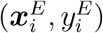 is a randomly chosen sample from an independent external dataset. Given the independence of ***X β*** and ***ϵ***, equation (3) can be expressed as

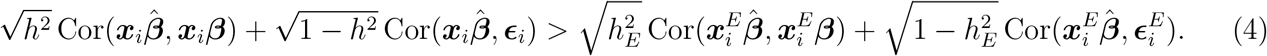

where 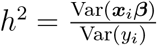 and 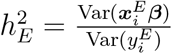 denote the heritability of the trait in the target and the external dataset respectively, and 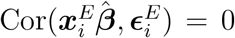 by definition. A sufficient condition for *no* overfitting is thus

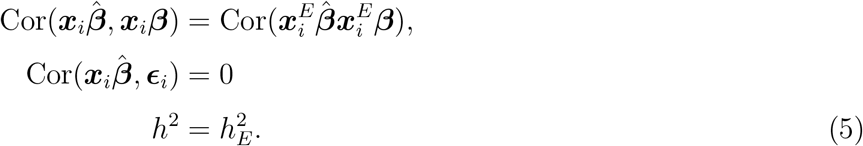

The fact that when the target data is used to calculate the summary statistics 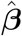, overfitting occurs, can be seen by considering a Directed Acyclic Graph (DAG), showing the relationship between 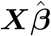 and ***X β*** (Fig 2(a)). (A DAG can be seen as a graphical representation of the probabilistic dependency of the different variables, and its interpretation is grounded in probability theory [30]. Two variables are ‘connected’ if a line can be traced through the graph connecting the two variables, except when a ‘collider’ is present along the path that connects the two. A ‘collider’ is a variable within a path where the two edges connecting it are both arrows pointing towards it, such as the variables ***y***, 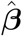, and 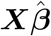, in Fig 2(a). Probabilistically, variables that are connected are expected to be dependent and correlated. Variables that are not connected are not dependent and thus not correlated.) We see that 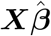 is connected to ***ϵ*** through ***y*** and thus expected to be correlated. Moreover, because in general we expect 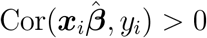 and Cor(*y_i_*, *ϵ*_*i*_) > 0, we expect 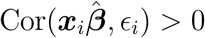, resulting in overfitting. In this article, we refer to this type of overfitting as OTD (Overfitting due to the overlap between the Target and the Discovery data).

**Figure 2:**
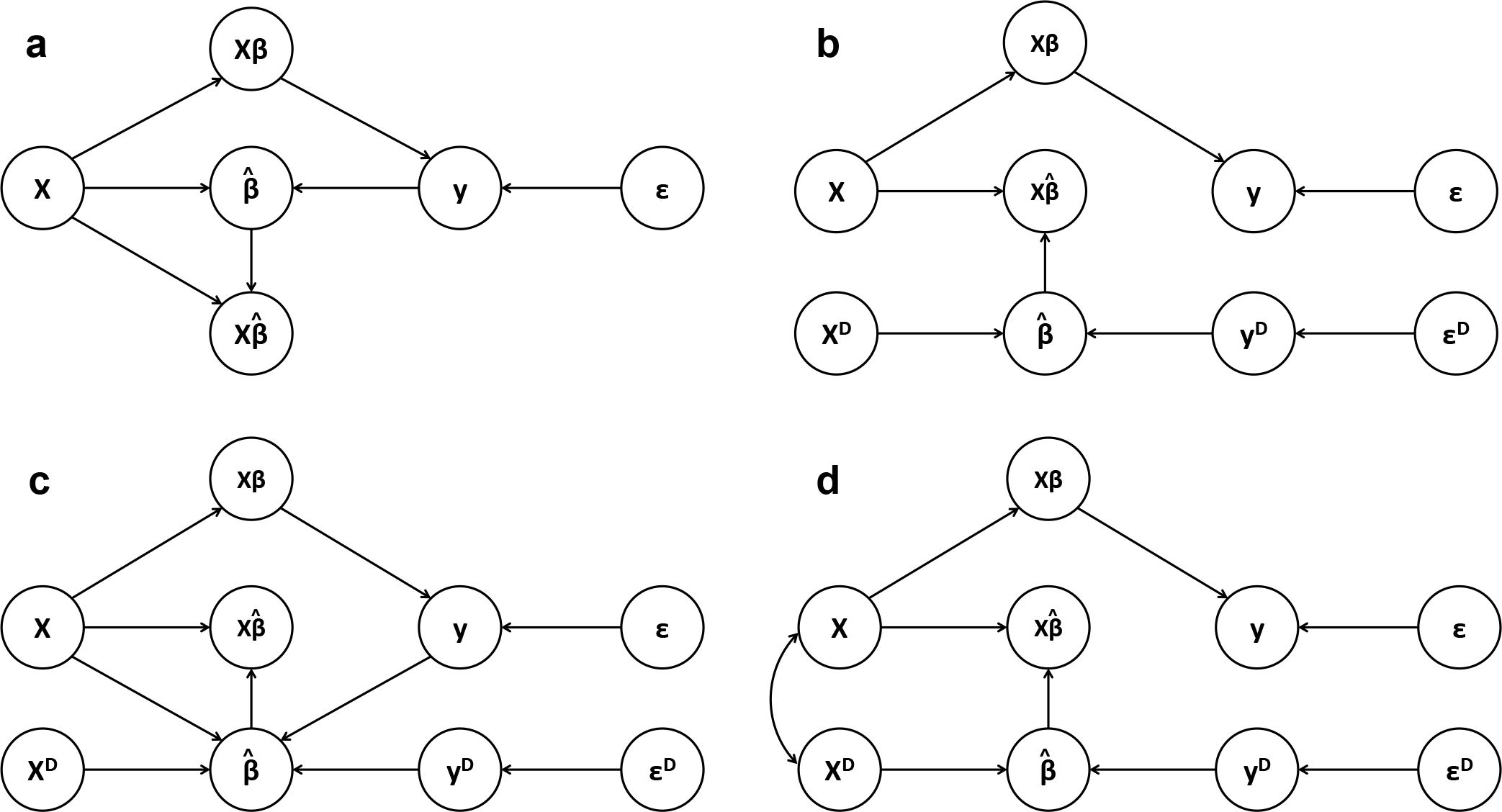
DAGs illustrating the relationship between the different variables in PGS estimation (a) when the target data is also used in the estimation of *β*, (b) when a separate discovery dataset (***X***^*D*^, ***y***^*D*^) is used, (c) when the target dataset is used in choosing the tuning paramter or the best 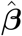 among a set of different 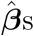, and (d) when the target dataset is genetically related to the discovery dataset.

In Fig 2(b) we see that if we use an external discovery dataset for estimating *β*, 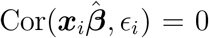, because the path between 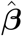 and ***y*** is broken. Moreover, if the external discovery sample ***x***^*D*^, ***x***^*E*^, and ***x*** are all drawn from the same population, 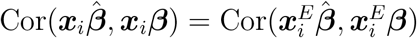 and overfitting is avoided.

A less appreciated kind of overfitting can be seen in Fig 2(c). Here, although the target dataset is not used for estimating ***β***, it is used for choosing a *p*-value threshold in the construction of PGS, as represented by the arrows pointing to 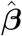 from ***X*** and ***y***. The fact that we generally choose the *p*-value threshold that maximizes the correlation between the PGS and the phenotype means that there is a Winner’s curse such that the apparent correlation between the PGS and the phenotype is higher than it would be in an external dataset. In this article we refer to overfitting due to the target data being used in validation OTV (Overfitting due to the overlap between the Target and the Validation data).

Finally, let us note that the inflation of correlation as cautioned by Wray *et al* [24] concerns not only the overlapping of samples. Rather, Wray *et al* [24] pointed out that inflation of correlation was likely if the target dataset were genetically related to the discovery dataset. We illustrate this situation in Fig 2(d), where correlations are expected between ***x*** and ***x***^*D*^. Here, although we still have 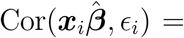 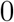 we cannot expect 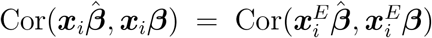, leading to overfitting. For the purposes of constructing PGS in large cohorts, however, this type of overfitting is of arguably less importance, since we are not interested in some external population ***x***^*E*^. In any case, accounting for differences in relatedness between the sample population and the general population at large is difficult and beyond the scope of this paper.

### Cross-prediction as a method to overcome overfitting due to the overlap of the target with the *discovery* data

As already noted above, overfitting can be avoided by breaking the path connecting ***y*** to 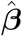. One way to do this in practice is to use an independent discovery dataset for estimating ***β*** (Fig 2(b)). When faced with a large target dataset which we also want to use as our discovery dataset, we can repeat this procedure in a cross-validation-like manner, i.e. we split the data into a number of folds, and repeatedly estimate 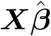 for the different folds, using the remaining folds for discovery. We call this procedure *crossprediction* (Figure 3(a)), to distinguish it from the more familiar procedure of cross-validation where fold-splitting is used only for choosing tuning parameters [31, 32, 33]. If external summary statistics are available, these can also be meta-analysed with those calculated from the discovery folds. To combine the PGS calculated in the different folds, we standardize them before stacking them together to form the final PGS. Standarizing and stacking them in this way will imply that the resulting stacked PGS represents the average correlation between the particular variable and the fold-specific PGS. Moreover, we prove that stacking the fold-specific PGS in this way preserves independence between individual elements of 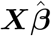 and ***ϵ*** and therefore does not introduce overfitting. Both of these proofs are presented in the Methods section.

### Split-validation as a method to overcome overfitting due to the overlap of the target with the *validation* data

In practical application of PGS, we do not simply have one PGS. Far more often, PGS are calculated for a range of *p*-value thresholds and the best one chosen. Letting 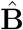 denote a matrix of coefficients where each column represent a vector of 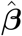 with different elements set to zero for different *p*-value thresholds, our estimated PGS is a matrix 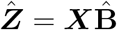 rather than a vector. This applies to each of the folds in cross-prediction, where for each fold *k* we have a matrix of standardized PGS 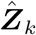. We need to choose the best column of 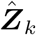 for each fold in order to form the final PGS, a step we refer to as *validation*. A common practice in PGS construction is to double up the target dataset as the validation dataset [13, 10, 34]. If put in the context of cross-prediction, this translates to performing validation and calculating the PGS within the target fold using the same data. However, as mentioned above, this can lead to overfitting, in particular OTV. Although the impact of this type of overfitting is commonly believed to be small, we illustrate its impact in the UK Biobank dataset by results from a simulation, whose details are given in the Methods section. Figure 4 shows the estimated correlations between multiple randomly generated (null) phenotypes and their PGSs calculated using cross-prediction. We see that although the phenotype was generated with no genetic component, when the target data doubled up as the validation data, the estimated correlations were inflated, compared to correlations with the phenotype in an external dataset. In Figure 5, we show this bias in terms of inflation in Type 1 error. When *p*-values between the estimated PGS and the phenotype are plotted against the expected distribution, there is a small but visible inflation in the statistics. Our strategy to overcome this is *split-validation*.In both Figures 4 and 5, we see that the method of *split validation* did not incur overfitting.

**Figure 3:**
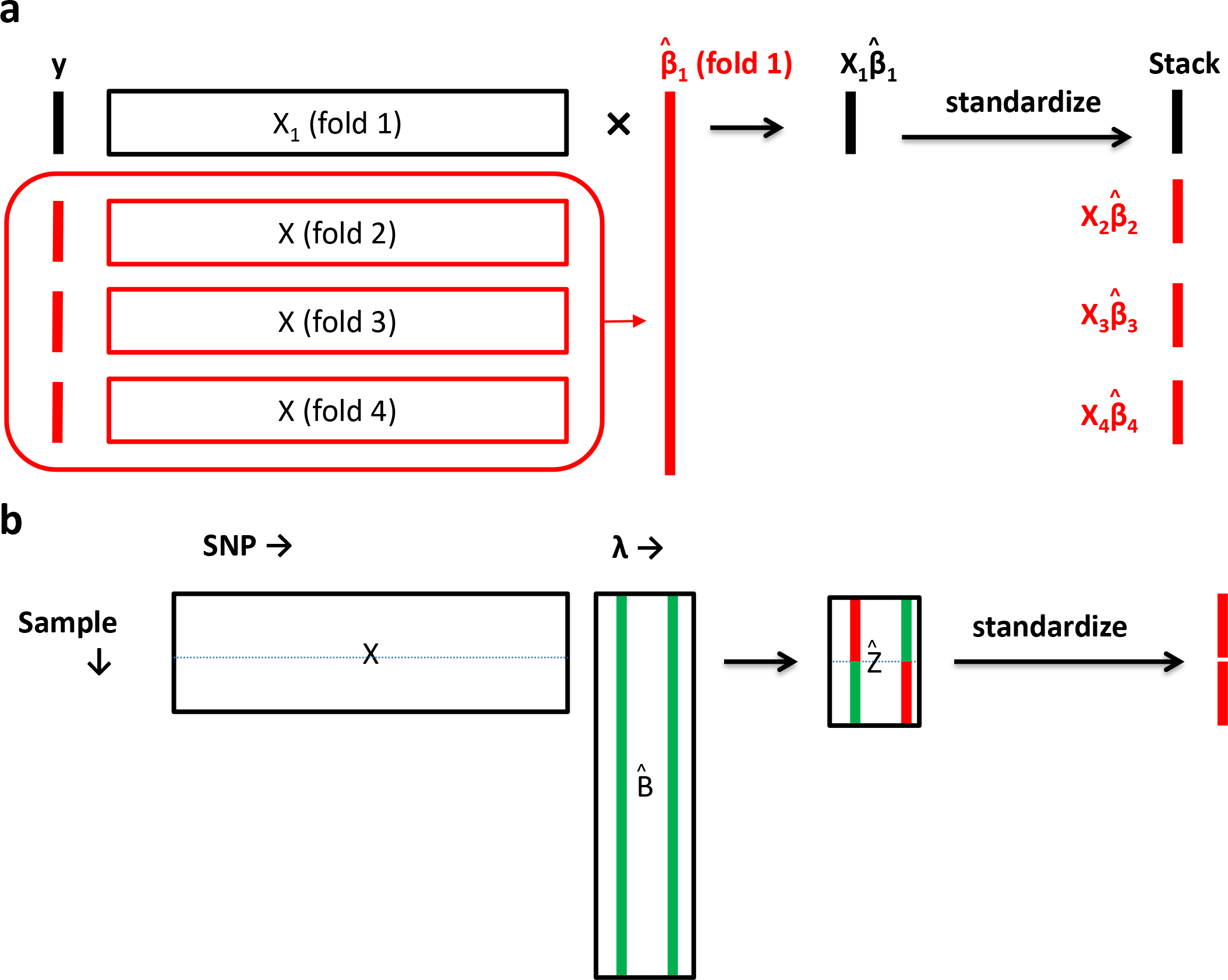
(a) Cross-prediction. The data (***X***, ***y***) is split into 4 folds. For fold 1, the coefficients 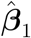 is estimated from folds 2,3, and 4. The estimated PGS 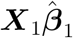 is standardized and stacked together to form the final PGS. (b) Split-validation. Let ***X*** deonte the genotype matrix, 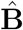 the matrix of coefficients, and λ indices the *p*-value threshold. Let 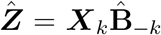 be the matrix of PGS calculated for the *k*^th^ fold. 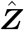 and ***X***_*k*_ are split into two halves. The green columns are the columns of 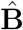 corresponding to the *p*-value threshold selected by validation. The red columns are the corresponding 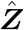 taken to form the PGS.Method for choosing *p*-value threshold

**Figure 4.**
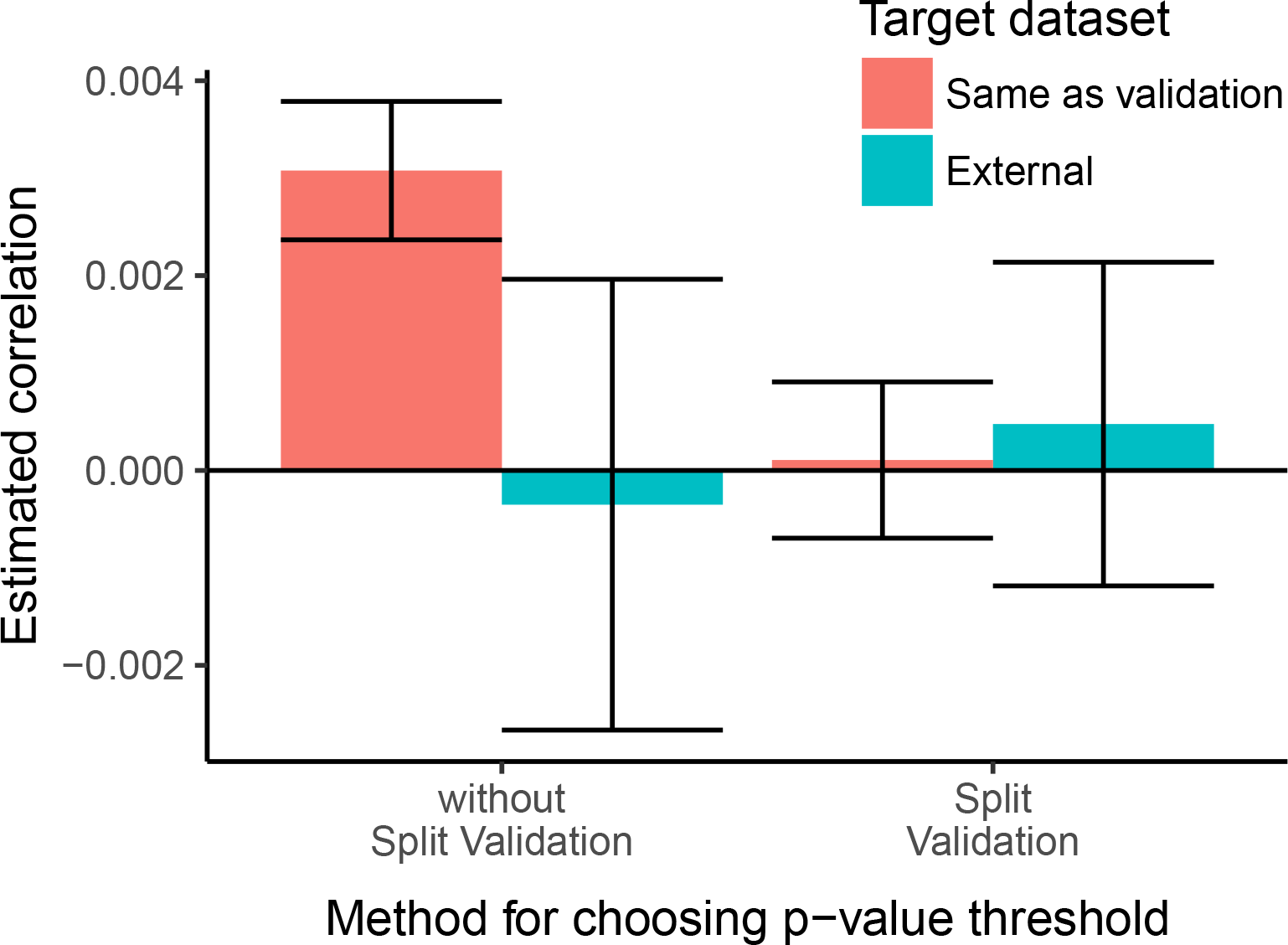
Barplots of mean estimated correlations between simulated null phenotypes (no genetic component) and the estimated PGSs, when the target dataset doubles up as the validation dataset, and when the target dataset is external. Error bars represent 95% confidence intervals.

The idea of split validation is similar to that of cross prediction. We first split the target dataset (or the target fold in cross-prediction) into two halves. We take turn to use each half for validation (i.e. the selection of the *p*-value threshold), and calculate the PGS in the other half using the *p*-value threshold selected in the other half and weights derived in the *discovery* dataset. A diagram illustrating split-validation is given in Figure 3(b).

**Figure 5:**
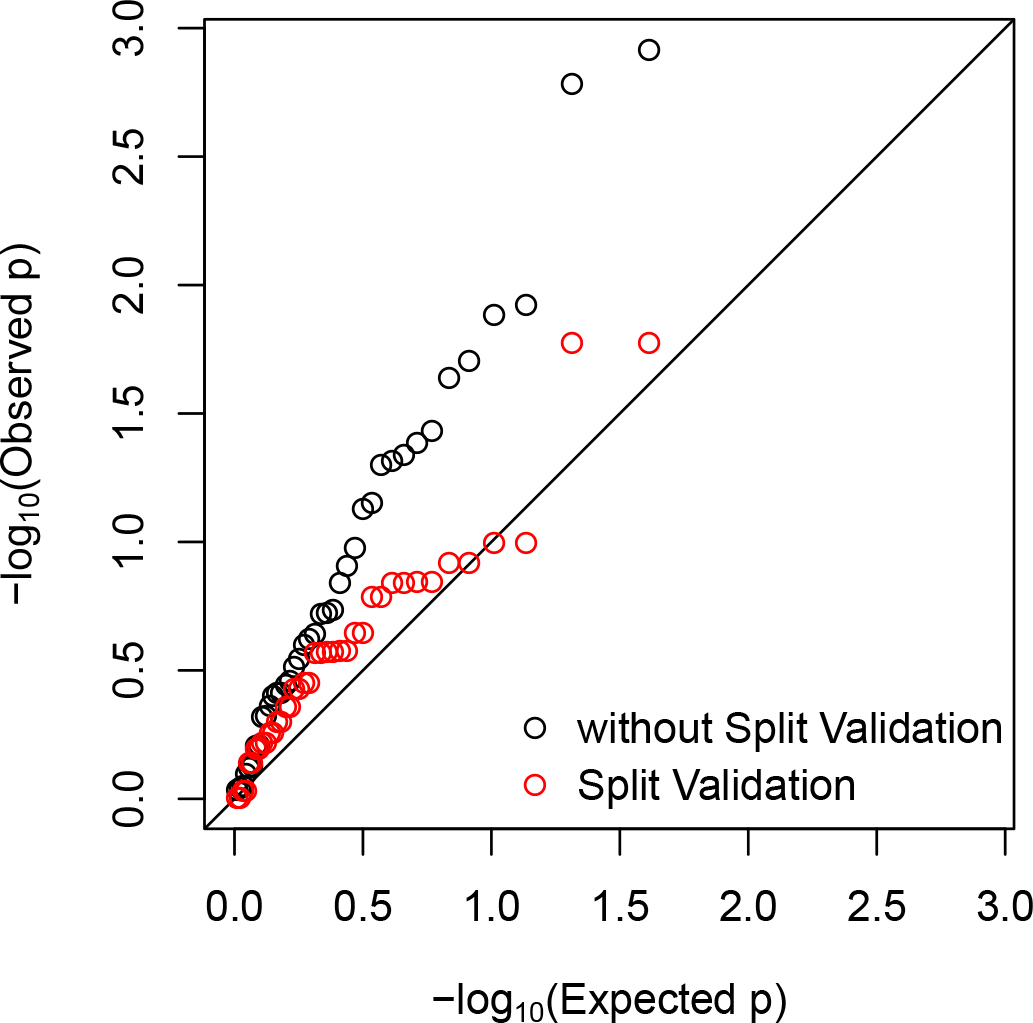
qq plot of *p*-values calculated for the relationship between the PGS and phenotype when the target data is also used for validation under a simulated null model.

**Figure 6:**
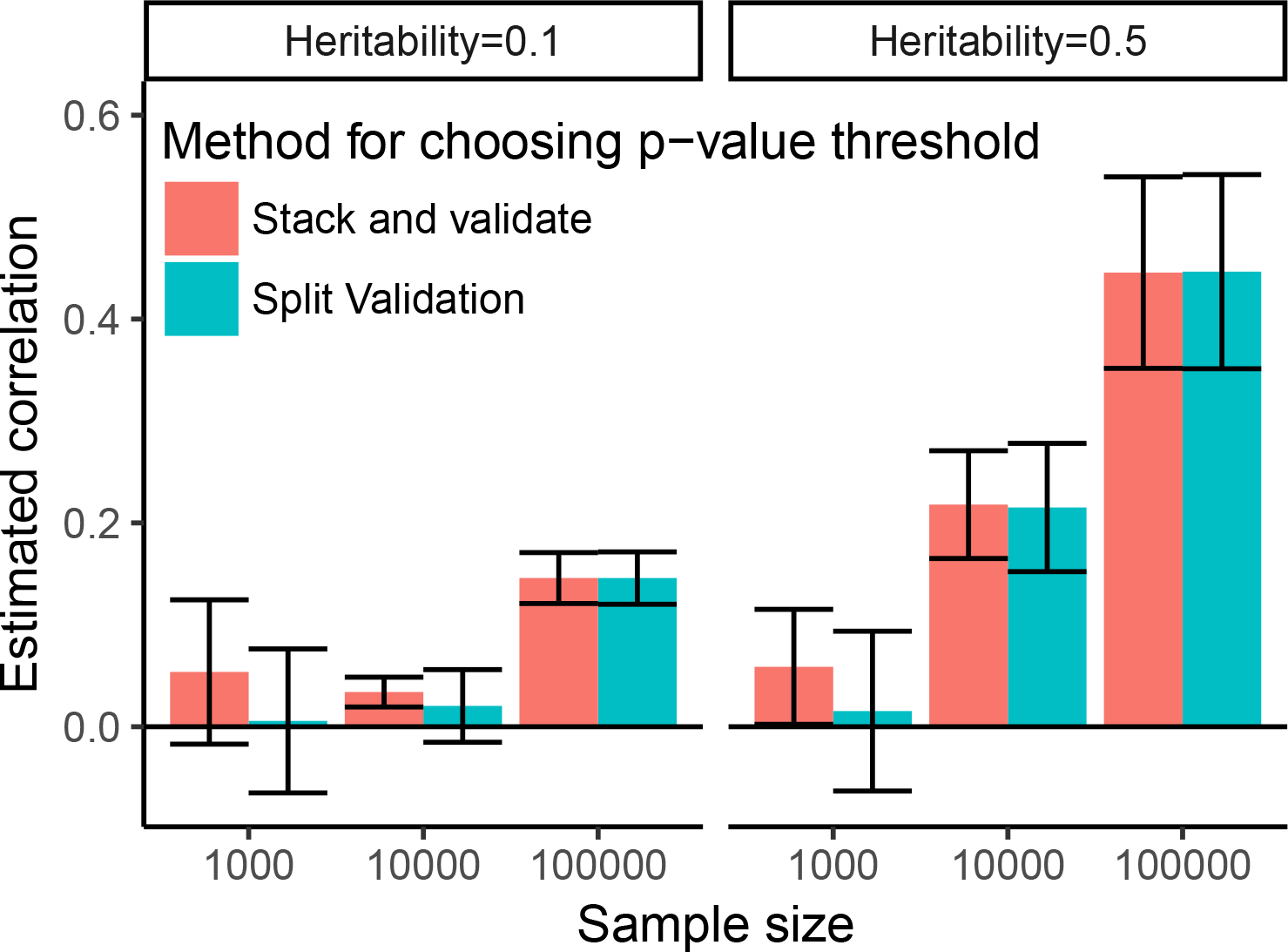
Comparison of split-validation vs “stack and validate” in calculating PGS. Mean and standard deviation of estimated correlation from 10 simulated phenotypes are plotted for each scenario.

### Sample size concerns

One possible concern with the cross-prediction + split-validation strategy is that instead of carrying out validation once, we carry out validation in multiple sub-samples within the target dataset, and this may reduce the power in choosing the best *p*-value threshold because of smaller samples. An alternative method that does not prevent OTV (but does prevent OTD) is to stack up the (standardized) PGS 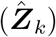 first (calculated for all *p*-value thresholds), and then validate them against the phenotype to choose the best *p*-value threshold. In Figure 6, we compare the performance of cross-prediction + split-validation vs the latter strategy (stack and validate) using sub-samples of the UK Biobank data and simulated phenotypes under two heritability scenarios (*h*^2^ = 0.1 and 0.5). 2,000 SNPs among 734,447 SNPs were assigned to be causal. It can be seen that when the sample size was 100,000, basically there were no difference in the predictive power of the PGS calculated using split-validation and stack and validate. When the sample size was smaller, we see that the predictive power of split-validation was reduced, particularly in the heritability=0.1 setting.

## Discussion

In this article, we show how cross-prediction, combined with split-validation, can be used to calculate PGS in large cohorts such as the UK Biobank. This can lead to a considerable increase in predictive power compared to using summary statistics alone. An overview of what constitutes *overfitting* is also given and it is shown that cross-prediction combined with split-validation overcomes both overfitting due to the target dataset overlapping with the discovery dataset (OTD) and with the validation dataset (OTV). The basic principle of the approach is the separation of the discovery, the validation, and the target subset of the dataset, and the combination of the resulting PGS from the different subsets through standardizing and stacking, which is shown preserve predicitve power and independence between the subsets.

One possible issue with this appraoch is that performing validation in different subsets and stacking the resulting PGS can reduce predictive power, compared to using the same data both for validation and prediction. However, it may be argued that with sample sizes of the magnitude of the UK Biobank, this is not an important issue.

In this article, we have not discussed overfitting due to other kinds of overfitting. In particular, we have not discussed possible overfitting due to the sample being related. Indeed it has been pointed out that the UK Biobank consists of a considerable number of second and third degree relatives [25]. This can lead to inflated estimates of the predictive accuracy of the PGS if estimates of *r*^2^ from the UK Biobank were extrapolated to the general population. On the other hand, we note that if our aim is to assess genetic correlation within the UK Biobank sample, then this type of overfitting is not relevant.

Usually genetic correlations can be assessed by examining the relationship between the PGS and various phenotypes. An important point to note is that overfitting can still occur when correlating different PGS calculated using the method of this paper. This is because in cross-prediction we try to keep the discovery and the target samples separate. However, when two PGS are both calculated using cross-prediction, their discovery samples can overlap, leading to overfitting.

We conclude with a number of suggestions for future work. First, depending on the number of folds use, a proportion of the sample is left out in the calculation of the summary statistics. It is unsure whether there can be a procedure that uses all data and also avoids OTD and OTV. Secondly, the current procedure is stochastic as the folds are randomly defined. The resulting PGS is also not a linear predictor in that it is not calculated as a linear combination of ***X***. Rather it is a mixture of different linear combinations. This has the disadvantage that theoretical properties of the PGS are less easily obtained. In principle, it is possible to find estimates of ***β*** such that when multiplied with ***X***, equals the CP PGS as calculated in our study. However, in our preliminary simulations, these estimates of ***β*** had very poor performance in external validation and we have not pursued this approach further. It is also possible in principle to extend this work further to the case where the number of folds used equals the sample size, such that we have a jackknife-like procedure for cross-prediction. This approach has not been studied. Thirdly, calculation of PGS using cross-prediction is currently time consuming for large cohorts and a large number of SNPs. In the simulations we have limited the number of SNPs to around 700,000. It may be possible to perform cross-prediction on a pre-selected set of SNPs using for example, informed pruning (clumping) [35]. However, if this set of SNPs were selected based on the entire dataset, overfitting would also arise, and future work is needed to minimize or avoid this bias.

## Methods

### PGS for six UK Biobank phenotypes

353,465 white British participants from the UK Biobank study were selected for these analyses. We used the genotype array of 734,447 SNPs, made available by the UK Biobank. The 6 phenotypes considered were height (ID=50), BMI (ID=21001), neuroticism (ID=20127), heart problems, taking medication for lowering cholesterol (ID=6153, 6177), and diabetes (ID=2443). Where multiple measurements were taken, the average was used. The variable “heart problems” was defined as a score from 0 to 3 based on the question “Vascular/heart problems diagnosed by doctor” (ID=6150), where 3 represents “Heart attack” or “Stroke”, 2 represents “Angina”, 1 represents “High blood pressure”, and 0 “None of the above”. The corresponding summary statistics were taken from the following studies: height[36], BMI[37], Neuroticism[38], Heart problems[39] (summary statistics for coronary artery disease), Medication for lowering cholesterol[40] (summary statistics for total cholesterol levels), Diabetes[41]. Only variants that were present in both the summary statistics and the UK Biobank genotype array were used for constructing PGS, both for the summary-statistics derived PGS and for cross-prediction. 5-fold cross-prediction was used with split-validation. All analyses, including the correlation between the phenotype and the PGS, were adjusted for the first 20 principal components and inferred gender. For Figure 1, selection of the *p*-value threshold for the summary-statistics only PGS was performed on an independent sample of 10,000 white British participants from the UK Biobank.

### Simulated phenotypes from the UK Biobank

For Figures 4 and 5, the same cohort of 353,465 white British participants from the UK Biobank was used. The phenotype was a simulated vector of 353,465 *N*(0,1) random variables and thus was completely independent of the genetic data. 5-fold cross-prediction was applied to compute the PGS. In the “without Split-Validation” scenario, the method of “stack and validate” was used (see Results section). The simulation was repeated 10 times and fold-specific correlations and *p*-values were plotted in the figures. The “external” target dataset was an independent dataset of 10,000 white British participants randomly selected from the UK biobank. No covariate adjustments were performed with these analyses.

For Figure 6, the linear model of (1) was used to generate the phenotype, with heritability, i.e. 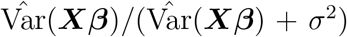 constrained to be either 0.1 or 0.5. Samples of size 1,000, 10,000, and 100,000 were randomly selected from the 353,465 white British participants.

### Details of PGS calculation

In all calculation of PGS in this paper, clumping and thresholding was used. First, summary statistics were clumped using the default settings in PLINK 1.9[42], where variants with an *R*^2^ of 0.2 or above within a 250kb region were “clumped” with the most significant SNP. *p*-value thresholds of 1*e*^−20^, 5*e*^−20^,1*e*^−19^, 5*e*^−19^,…, 0.001, 0.005, 0.01, 0.02, 0.03,…, 0.99,1 were used. Clumping and *p*-value thresholding was performed independently for each fold in cross-prediction.

### Computation

An R package (`crosspred`) has been written to perform cross-prediction and split-validation, and is available on https://github.com/tshmak/crosspred/blob/master/CrossPrediction.md. The package is designed to be a wrapper around the package `lassosum` [43]. Although clumping and *p*-value thresholding was used throughout this paper to calculate PGS (as it is the more widely used method), in principle, it is possible and even preferable to use `lassosum` instead, which can lead to better predictive power.

#### Proof: standardizing PGS within fold before stacking approximates the average correlation of the PGS with another variable

Let 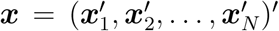 denote a stacked column of PGS, and ***y*** a column of phenotype. Further assume ***x*** is standardized within fold, such that **1′x**_*k*_ = **0** and 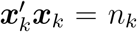, and that ***y*** is standardized such that ***1′y*** = **0** and **y′y** = n = ∑_*k*_ *n*_*k*_ without loss of generality. The correlation of ***x*** with ***y*** is ***x***–***y***/*n*. Let the standard deviation of ***y*** within fold *k* be 1/*s*_*k*_. We have

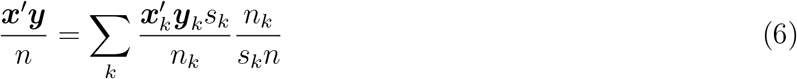

where 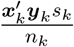 is the fold-specific correlation. Thus, 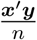 is a weighted average of the fold-specific correlation with weights 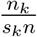. In general *s*_*k*_ approximates 1, such that the weights are approximately optimal.

#### Proof that 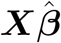 remain independent of with *ϵ* after stacking

As in the main text, we assume that 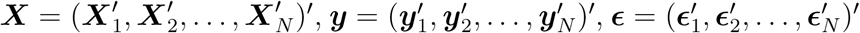. Denote 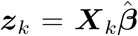 From Figure 2(b), we establish that *z*_*k*_ is independent of ***ϵ*** if 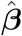 is derived from a different fold from ***z***_*k*_. It follows that the *i*^th^ element of ***z***_*k*_, denoted *z*_*ki*_ is independent of the *i*^th^ element of ***ϵ***, within a particular fold 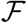. In notation:

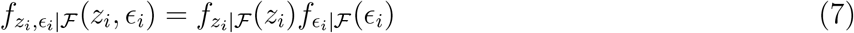

*Proof:* 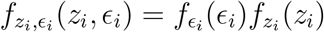.

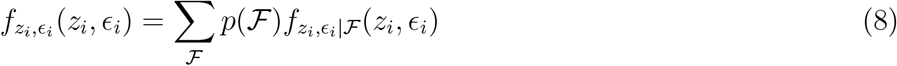

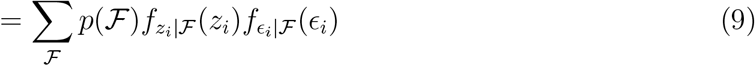

Now, because *ϵ*_*i*_ are assumed *i.i.d*. regardless of fold, we have

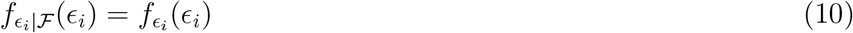

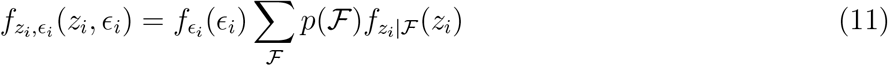

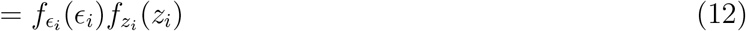

completing the proof.

## Acknowledgement

This research has been conducted using the UK Biobank Resource (project ID 23079, 28732)

